# Absence of estrogen leads to defects in spermatogenesis and increased semen abnormalities in male rabbits

**DOI:** 10.1101/2022.10.10.511525

**Authors:** Aurélie Dewaele, Emilie Dujardin, Marjolaine André, Audrey Albina, Hélène Jammes, Frank Giton, Eli Sellem, Geneviève Jolivet, Eric Pailhoux, Maëlle Pannetier

**Author notes:** Both authors contribute equally to this work (Eric Pailhoux, Maëlle Pannetier). (A.D.), (E.D.), (M.A.), (A.A.), (H.J.), (G.J.), (E.P.), (M.P.), (F.G.), (E.S.). Correspondence: (E.P.); (M.P.).

## Abstract

Estrogens are steroid hormones produced by the aromatization of androgens by the aromatase enzyme, encoded by the *CYP19A1* gene. Although generally referred to as “female sex hormones”, estrogen is also produced in the adult testes of many mammals, including humans. To better understand the function of estrogens in the male, we used the rabbit model which is an important biomedical model. First, the expression of *CYP19A1* was localized, demonstrating that testicular estrogens are produced by meiotic germ cells inside the seminiferous tubules. Next, the cells expressing ESR1 and ESR2 were identified, showing that estrogens could exert their function on post-meiotic germ cells in the tubules and play a role during sperm maturation, since ESR1 and ESR2 were detected in the *cauda* epididymis. Then, CRISPR/Cas9 *CYP19A1*^−/−^ genetically modified rabbits were analyzed. *CYP19A1*^−/−^ males showed decreased fertility with lower sperm count associated with hypo-spermatogenesis and lower spermatid number. Germ/sperm cell DNA methylation was unchanged, while sperm parameters were affected as *CYP19A1*^−/−^ males exhibited reduced sperm motility associated with increased flagellar defects. In conclusion, testicular estrogens could be involved in the spermatocyte-spermatid transition in the testis, and in the acquisition of sperm motility in the epididymis.

## 1. Introduction

Sex steroids are key reproductive system hormones in both sexes. Estrogens have always been considered the female sex steroid hormones and androgens as their male counterparts. This simplistic assessment remains accurate in different species of vertebrate and for several developmental pathways such as ovarian development in non-mammalian species in which estrogen plays a key role [1–5] or in the differentiation of internal and external male genitalia of mammals for androgens [6]. However, it is clear, since many decades, that the situation is more complex; on the one hand because the synthesis of estrogens is made from androgens thus implying the presence, at least transitory, of androgens in females, and on the other hand because estrogens are also produced by the testes of mammals where their roles remain to be elucidated (for review see [7]).

Cytochrome P450 aromatase, encoded by the *CYP19A1* gene, is responsible for the irreversible conversion of androgens to estrogens. This enzyme is expressed in the adult testes in mammals, but its cellular localization is highly variable depending on the species and the laboratory of analyses (for review see [8]). Initially, the aromatase expression was described in Leydig cells in rats [9], pigs [10], stallions [11] or humans [12]. Its expression in Sertoli cells was also observed in immature rat testes [13] and aromatase was finally described in meiotic and post-meiotic germ cells of mice [14], rats [9] and humans [15]. Some studies even detected its expression in spermatozoa in pigs [16] and humans [15].

To promote their actions, estrogens are known to use two nuclear receptors ERα/ESR1 and ERβ/ESR2, resulting in genomic effects; and a G-protein-coupled seven-transmembrane receptor (GPER, G-Protein Estrogen coupled Receptor) causing rapid non-genomic effects. On the base of the literature, ERs expression can be detected in all testicular cell types although the results often differ between species and studies (reviewed in details by [17]). For example, in the human testis, ESR1 expression has been described in Leydig cells [18], or in spermatogonia, spermatocytes and round spermatids [19], or in Leydig, Sertoli and germ cells [20]. This variability of results can be explained by the existence of ER variants (spliced isoforms [21]), other proteins that share homology with classical ER, or by the methodologies and antibodies used.

The importance of estrogen signaling in the male fertility has been indicated by the adverse effects of estrogen-like compounds and their interaction with estrogen receptors, which have been shown to cause pathologies. In rats, nuclear receptor overstimulation experiments revealed the presumed role of estrogens in spermatogenesis. Treatment with an ESR1 agonist impaired the formation of elongated spermatids, while administration of an ESR2 agonist induced spermatocyte apoptosis and spermiation failure both leading to reduced sperm count [22]. In addition, overexposure to estrogen during spermatogenesis resulted in epigenetic defects in sperm, such as increased histone retention (ESR1 agonist) and decreased DNA methylation (ESR2 agonist) [23,24].

On the other hand, gene modification experiments carried out in mice tend to show that estrogens, in the testes and male genital tract, act mainly via the ESR1 receptor. Indeed, data on ESR2 functions in the male tract are still controversial in mice, since some showed normal *Esr2^−/−^* male fertility [25], while others described infertility of unknown origin [26]. In addition, mice deficient for the membrane receptor (*Gper^−/−^*) are fertile and show no particular phenotype [27]. On the contrary, a complete infertility was described in male *Esr1^−/−^* [28]. Early in their reproductive life, *Esr1*^−/−^ males showed testes with disorganized seminiferous epithelium and dilated lumen. While sperm counts were normal in *Esr1*^−/−^ males, spermatozoa presented reduced motility (flagellar defects)[28–30] and were ineffective in *in vitro* fertilization (premature acrosomal reaction in mutants) [28,30]. The latter phenotypes appeared to be related to epididymal dysfunctions, and alterations of the epididymal fluid milieu were observed in *Esr1*^−/−^ mice [29].

Genetic modifications or mutations affecting estrogen production have also been reported. In mice, in the absence of aromatase in males (*Cyp19a1^−/−^* or ArKO), normal testicular morphology was observed up to 14 weeks, with no signs of infertility. Then a progressive alteration of spermiogenesis was reported, leading to an increase in apoptosis of round spermatids and degeneration of the seminiferous epithelium: the ArKO mice became infertile with advancing age [31,32]. In humans, aromatase mutations are extremely rare conditions. In these patients, there are no consistent findings regarding the testicular phenotype (review in [33]). Nevertheless, when semen collection could be done, oligo-azoospermia and reduced sperm motility were observed [34,35].

The rabbit is an important biomedical model that could help to better understand the function of this testicular estrogen production for spermatogenesis. Thus, we first described *CYP19A1*, *ESR1* and *ESR2* expression in the testis and epididymis of adult rabbits. We showed that estrogens are exclusively produced by germ cells, mainly pachytene spermatocytes. Both *ESR1* and *ESR2* were expressed by round spermatids. Additionally, these receptors were detected in the epididymis, mainly the *cauda*, where estrogen could be measured. Then, taking advantage of the *CYP19A1* mutant rabbit model created by our laboratory [36], we investigated the effects of estrogen deprivation on testes and sperm production in this species. First, a slight decrease in fertility was observed in homozygous mutant males. Then, abnormalities of the seminiferous epithelium were observed, which were related to impaired spermatogenesis and led to a lower sperm count. Finally, sperm motility was affected and sperm morphological abnormalities were increased in mutant males suffering from estrogen deprivation.

## 2. Materials and Methods

### 2.1. Animals

New Zealand rabbits (NZ1777, Hypharm, Rousssay, France) were bred at the SAAJ rabbit facility (Jouy-en-Josas, France). All experiments were performed with the approval of the French Ministry MENESR (accreditation number APAFIS#6775-2016091911407980 vI) following the recommendation given by the local committee for ethic in animal experimentation (COMETHEA, Jouy-en-Josas). All researchers working directly with the animals possessed an animal experimentation license delivered by the French veterinary services. Three independent lines of *CYP19A1* mutant rabbits have been generated [36] and two of them have been used in this study since no phenotypical differences have been observed.

From sexual maturity (6 months), heterozygous *CYP19A1^+/−^* and homozygous *CYP19A1^−/−^* males were mated with heterozygous *CYP19A1^+/−^* females, while control males and females were mated together. The number of mating with or without birth per male, as well as the number of pups per litter was monitored.

### 2.2. Histological and immunohistological analyses

Immediately after sampling, pieces of adult testes were immersed in Bouin’s fixative or paraformaldehyde (4% PFA in PBS 1x), fixed for 72 hours then paraffin embedded. Microtome sections of 5μm thickness were processed. Periodic Acid Schiff (PAS) colorations were performed by the @Bridge platform (INRAE, Jouy-en-Josas) using an automatic Varistain Slide Stainer (Thermo Fisher Scientific).

*In situ* hybridization (ISH) was performed using the RNAscope ISH methodology (ACD, Bio-Techne SAS, Rennes, France) as previously described [36]. *CYP19A1*, *ESR1* and *ESR2* probes that have been used were those published previously [36]. Hybridization was performed on 5μm sections from PFA fixed tissue using labelling kits (RNAscope 2.5HD assay-brown (conjugated to horse radish peroxidase)) as recommended by the manufacturer. Hybridization was considered as positive when at least one dot was observed in one cell. All colored sections (visible) were scanned using a 3DHISTECH panoramic scanner at the @Bridge platform (INRAE, Jouy-en-Josas).

Immunofluorescence was performed on rehydrated sections, where epitope retrieval was performed with citrate-based unmasking solution in a pressure cooker. DNA was then denatured 15 min with HCl 2N, and sections were permeabilized by incubation with 0.5% Triton, 1% BSA for 1h30. After an overnight incubation at 4°C with the primary antibodies (anti-5mC, Eurogentec, ref BI-MECY-0100, 1/500; anti-5hmC, Active Motif, ref 39569, 1/500), and a 1-hour incubation at room temperature with secondary antibodies (Dylight 488 anti-mouse, KPL, ref 072-03-18-06, 1/200; anti-rabbit AlexaFluor 488, Life Technologies, ref A21441, 1/200), slides were mounted in Vectashield mounting medium (Vector) containing DAPI and images were acquired with a DP50 CCD camera (Olympus).

### 2.3. RNA extraction and RT-qPCR analyses

The testis and epididymis (head and cauda) from adult rabbits were collected and immediately frozen at −80°C. Total RNA from each sample was extracted using the RNeasy^®^ MicroKit (Qiagen, France). Quantitative PCR was performed on reverse transcribed RNAs (High Capacity Reverse cDNA Transcription kit with the included set of random primers, Applied Biosystems, ThermoFisher, France). Based on the output of the GeNorm program, we used *H2AFX*, *YWHAZ* and *SF1* (*Splicing Factor 1*) as the reference genes for this study (Table 1). The results were analyzed with qBase Software [37].

**Table 1.**
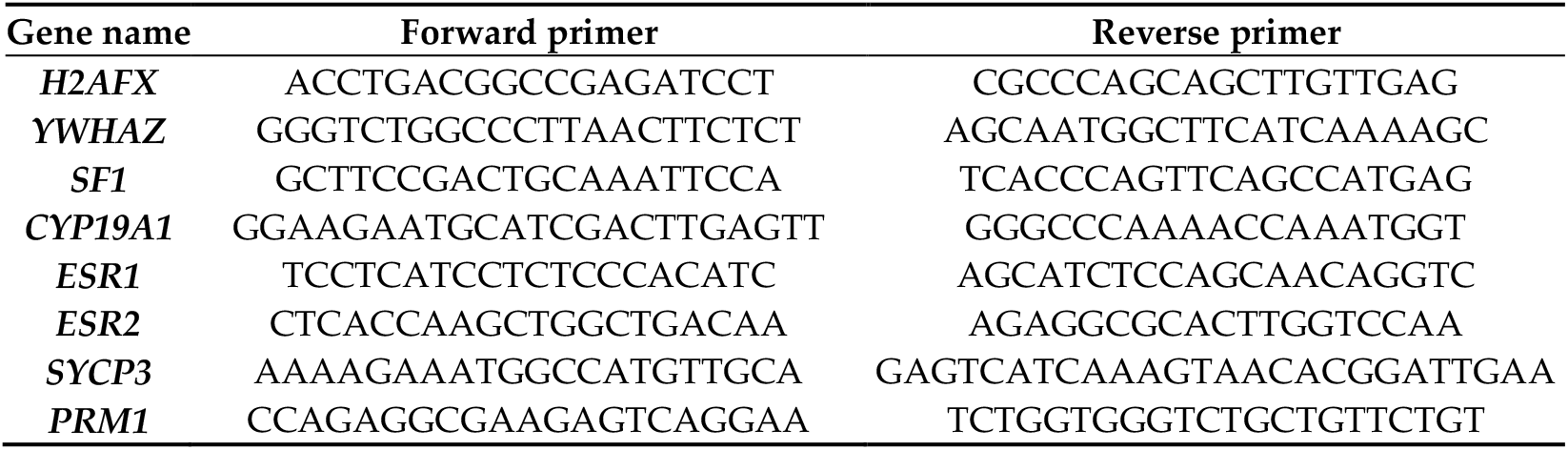
Primers used by RT-qPCR.

### 2.4. Measurement of estradiol, testosterone and DHEA hormone levels in testis and epididymis of adult rabbits

Estradiol, testosterone and DHEA were assayed by GC/MS according to the protocol described by [38] with modifications [39]. Sample extraction and purification, derivatization and determination of steroid levels in testes and epididymides from adult rabbits are described in [36] or can be provided upon request.

### 2.5. Semen collection and sperm parameter analyses (CASA)

Semen of rabbits from the different genotypes were collected using a specially designed artificial vagina. Two successive sampling have been made when possible for each animal. The ejaculate volume was estimated by pipetting and sperm was then immediately diluted in GALAP media (IMV Technologies) which was specifically designed for the conservation of rabbit semen. Each diluted samples was incubated 10 min at 37°C before analyzing sperm parameters using a CASA Hamilton Thorne IVOS II device with the x10 objective.

### 2.6. Luminometric Methylation Assay (LUMA)

DNA extraction from sperm samples was performed as described in [40]. Global DNA methylation levels were quantified using LUMA [41]. Briefly, 1 μg of genomic DNA was cleaved using the isoschizomers HpaII (methylation sensitive) and MspI (non-methylation-sensitive) in two separate reactions and in the presence of EcoRI to standardize for DNA amounts (New England Biolabs). The protruding ends were then used as templates for pyrosequencing with the Pyromark Q24 device and Pyromark Gold Q96 reagents (Qiagen). The luminometric signals produced by either the sequential incorporation of C and G nucleotides (reflecting the number of CCGG sites digested by HpaII or MspI) or the sequential incorporation of A and T nucleotides (reflecting the number of AATT sites digested by EcoRI), were then quantified using Pyromark Q24 software. Each sample was assayed in duplicate.

The global methylation percentage per sample was then calculated as follows: Methylation% = [100−(Average signal obtained with HpaII after EcoRI normalization/Average signal obtained with MspI after EcoRI normalization)] * 100

### 2.7 Statistics

The statistical analyses were performed using the GraphPad Prism 7 Software (GraphPad Software Inc., La Jolla, CA, USA). Because of the small number of samples in groups, comparisons between values were made by the Mann-Whitney test for non-parametric values. A probability lower than 0.05 was required for significance.

## 3. Results

### 3.1. Localization of CYP19A1/aromatase and ESRs expression in the rabbit testis and epididymis

To decipher which testicular cell type expressed the *CYP19A1* gene and thus, were able to produce estrogens, we carried on *in situ* hybridization by using the RNAscope technology, giving clear and reproducible results on rabbit gonads [36] (Figure 1). Aromatase expression was detected in germ cells exclusively. These cells which were closed to the basal lamina, presented a large nucleus and seemed to correspond to pachytene spermatocytes (Figure 1A). A few labelling was also detected in the round spermatids lying in close proximity of the pachytene cells, suggesting the persistence of some aromatase transcripts in post-meiotic germ cells. In addition, to determine which cell types could respond to estrogens in the seminiferous compartment, probes corresponding to estrogen receptors *ESR1* and *ESR2* were used. The expression of both estrogen receptors was attested into the seminiferous tubules and was restricted to the round spermatids, with a stronger labelling for *ESR1* (Figure 1C).

**Figure 1.**
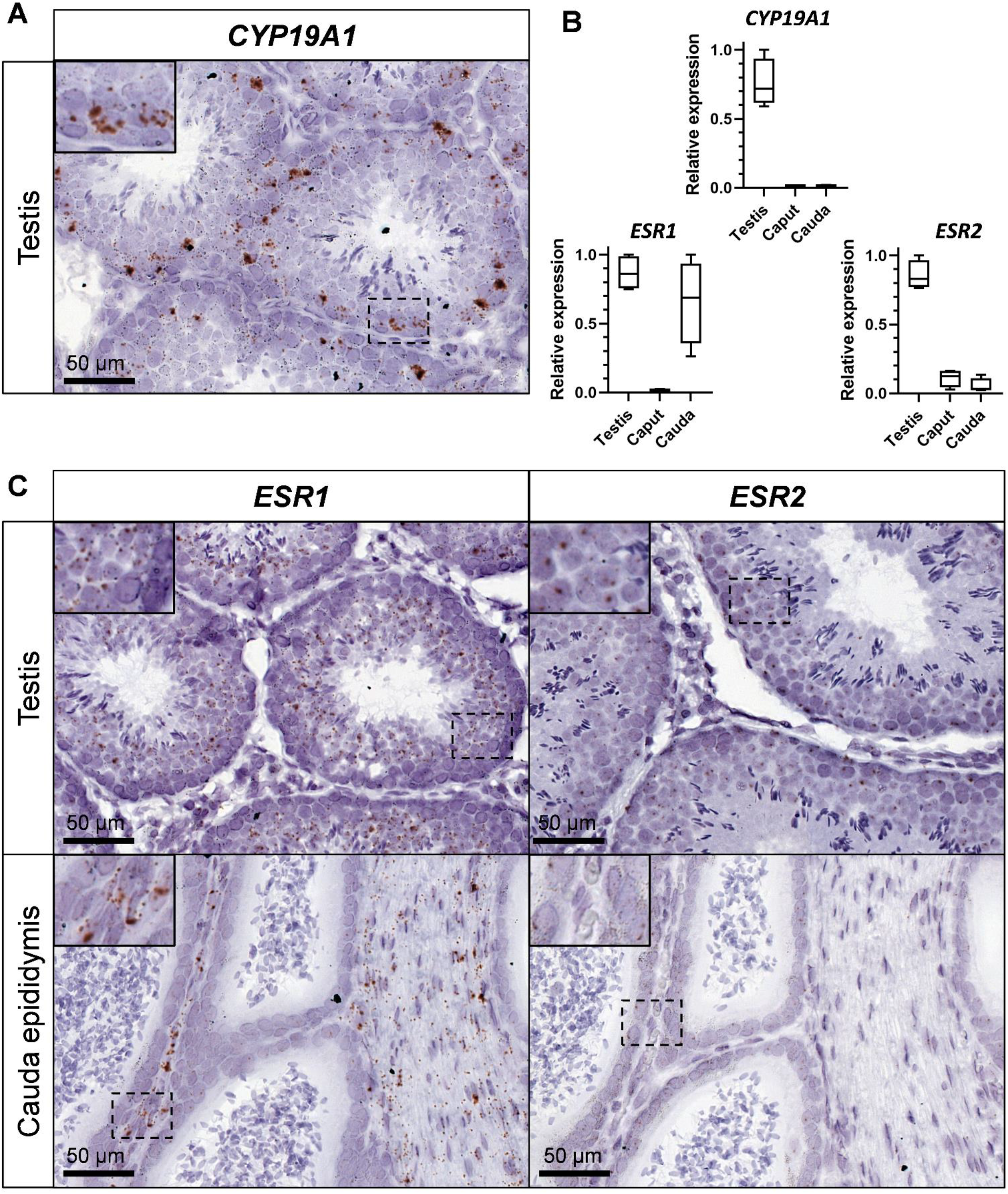
*CYP19A1* and estrogen receptors expression in the testis and epididymis of adult rabbits. (**A**) Location of *CYP19A1* expression by *in situ* hybridization (RNAscope technology) in the adult testis. (**B**) Relative expression levels of *CYP19A1, ESR1* and *ESR2* analyzed by RT-qPCR in the adult testis and epididymis (caput and cauda). Testis n=4; Epididymis n=4. (**C**) Location of *ESR1* and *ESR2* expression by *in situ* hybridization (RNAscope technology) in the testis and the cauda epididymis.

The expression level of both aromatase and estrogen receptors transcripts was studied and compared within testes and epididymides (*caput* and *cauda*) by RT-qPCR (Figure 1B). Although *CYP19A1* expression was restricted to the testis, *ESR1* expression was strongly detected in the testis and in the tail of the epididymis (*cauda*). *ESR2* expression was also observed in testes, and was faintly detected in both the head and the tail of the epididymis. However, *in situ* hybridization failed to detect *ESR2* (or *ESR1*) expression in the *caput* epididymis (Figure 1C). On the contrary, strong staining was obtained in the mesenchyma with *ESR1* probe and a faint staining for *ESR2* in the epithelial cells of the *cauda* epididymis.

### 3.2. CYP19A1 gene-targeting in rabbits efficiently suppresses testicular estrogen secretion

We have previously established three strains of genetically modified rabbits carrying deletions of exon 2 (including the initiator ATG codon) of the *CYP19A1* gene. These rabbit strains were initially created to evaluate the role of fetal estrogens produced by early developing ovaries (i.e.: before meiosis initiation in the germinal lineage) in a non-rodent mammalian species [36]. To confirm that knocking out *CYP19A1* in male really led to testicular estrogen deprivation, we measured estrogen concentrations in *CYP19A1^−/−^* testes in comparison with wild type ones (Figure 2A). As expected, consistent with estrogen assays performed for their female counterparts [36], no testicular estrogen remained detectable in mutant gonads compared to wild type ones where the median 17β-Estradiol value was about 55 pg/g of tissue. Even if testicular estrogen production was abolished, testosterone and dehydroepiandrosterone (DHEA) concentrations remained similar between mutant and wild type testes with respectively, around 9 ng/g and 40 ng/g (median values) in each condition (Figure 2A). These results confirm that the deletions engineered on *CYP19A1* exon 2 in the three rabbit lines efficiently suppressed aromatase activity and estrogen secretion as previously demonstrated in females [36].

**Figure 2.**
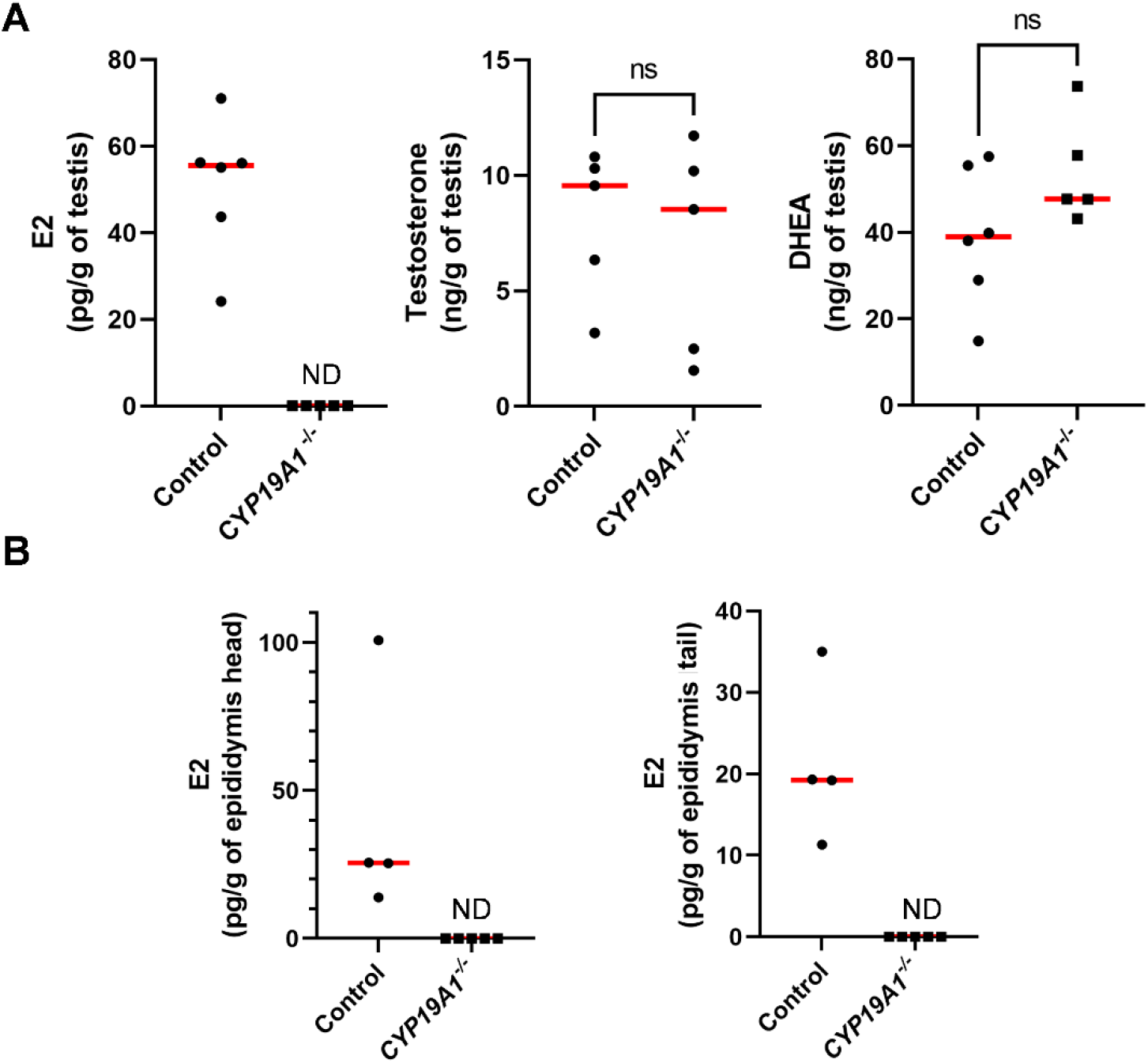
Steroid levels in the testis and epididymis in adults. (**A**) Dosage of 17β-estradiol (E2), testosterone and DHEA concentrations in control and *CYP19A1^−/−^* adult testes by GC/MS. (**B**) Dosage of 17β-estradiol (E2) in the head and the tail of epididymis. Control n = 4 to 6; *CYP19A1*^−/−^ n = 5. The median is shown in red. ND: Non Detected. Mann-Whitney test: * pValue<0.05. ns: non significant.

We have previously shown that estrogens are produced in the seminiferous tubules. Due to the presence of the blood-testis barrier, estrogens were expected to circulate through the efferent ducts and epididymides. 17β-Estradiol levels were thus measured in the epididymides, showing 25 pg/g and 20 pg/g as median values in the head and the tail of the epididymis respectively (Figure 2B). No estrogen could be detected in *CYP19A1* homozygous mutant epididymides.

### 3.3. CYP19A1 knockout male rabbits show subfertility parameters

During our previous study on *CYP19A1*^−/−^ fetal ovary [36], spanning on 8 years, *CYP19A1* genetically modified male rabbits from three different strains have been mated with heterozygous females (*CYP19A1*^+/−^) to expand the lines and produce biological materials. Either heterozygous carrier males (XY *CYP19A1*^+/−^) or homozygous mutant males (XY *CYP19A1*^−/−^) were used. Interestingly, a slight subfertility of homozygous mutant males compared to heterozygous carrier ones was observed on two recorded parameters. First, the percentage of mating without birth was increased when using homozygous mutant males (58.9%) compared to heterozygous carrier males (32.3%) or control rabbits (35.5%) (Figure 3A). Secondly, when mating was successful, the recorded litter size was statistically reduced, as the number of pups per litter dropped from 7.9 (± 3.3) to 6 (± 2.8) by using heterozygous or homozygous mutant males respectively (Figure 3B and 3C).

**Figure 3.**
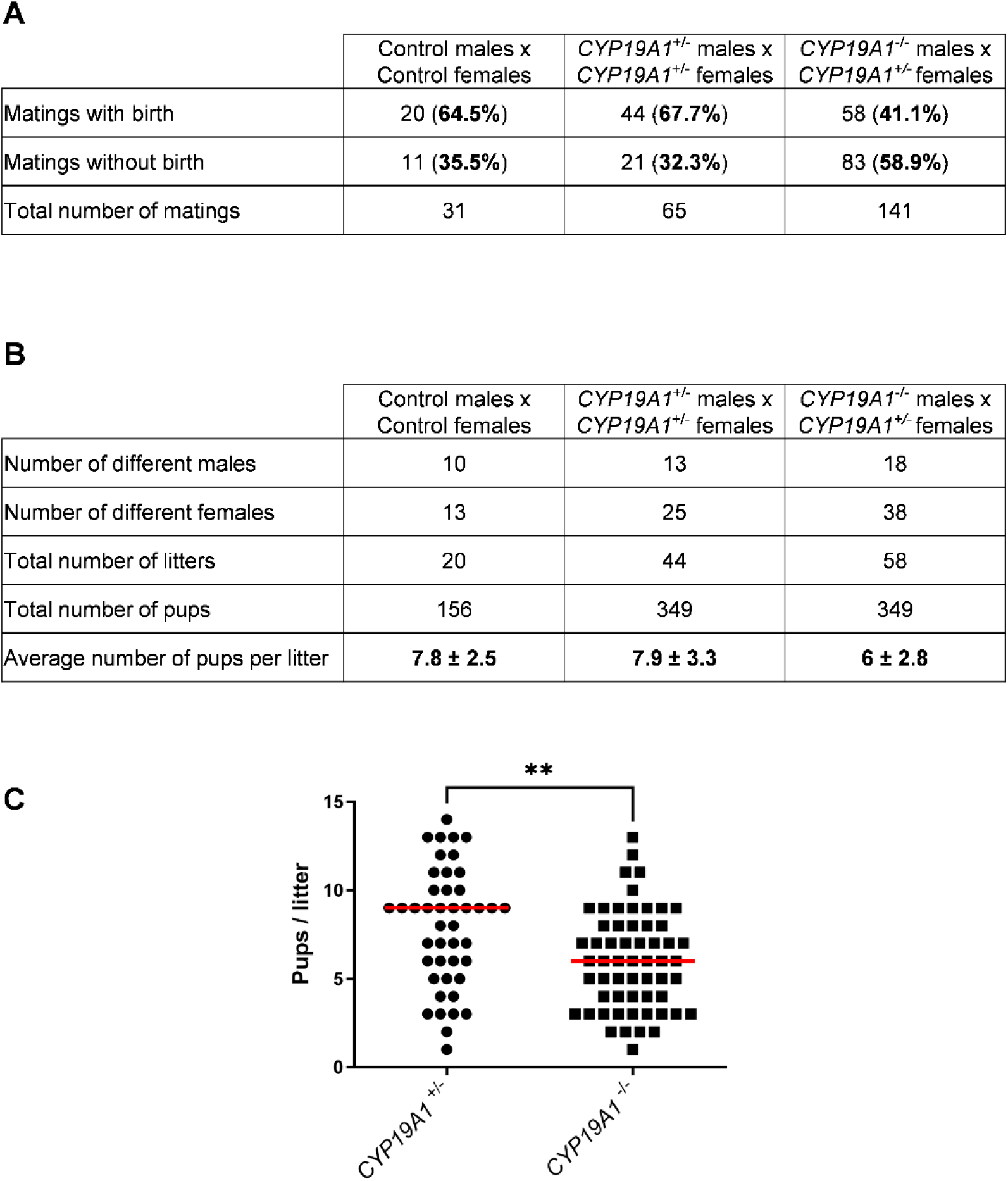
Effect of the *CYP19A1* gene knock-out on the male fertility. (**A**) Number of successful and unsuccessful mating, depending on the genotype of the parents. (**B**) Number of different males and females used and pups per litter depending on the genotype. (**C**) Number of pups per litter obtained by crossing heterozygous females with heterozygous (*CYP19A1*^+/−^) or homozygous (*CYP19A1*^−/−^) males. The median is shown in red. Mann-Whitney test: ** p<0.01

### 3.4. Absence of testicular estrogens leads to spermatogenesis defects

The effects of estrogen deficiency on testicular morphogenesis and function were evaluated in 2- to 3-year-old rabbits. Histological analyses of the testes showed some abnormal seminiferous tubules in the homozygous mutants (Figure 4A-F). These have been found clustered in a few testicular lobules or scattered through the testis (Figure 4C and D and Figure 4E and F respectively). In these abnormal tubules, the thickness of the seminiferous epithelium was reduced and the lumen appeared larger: a drastic decrease of the spermatid layer was observed.

**Figure 4.**
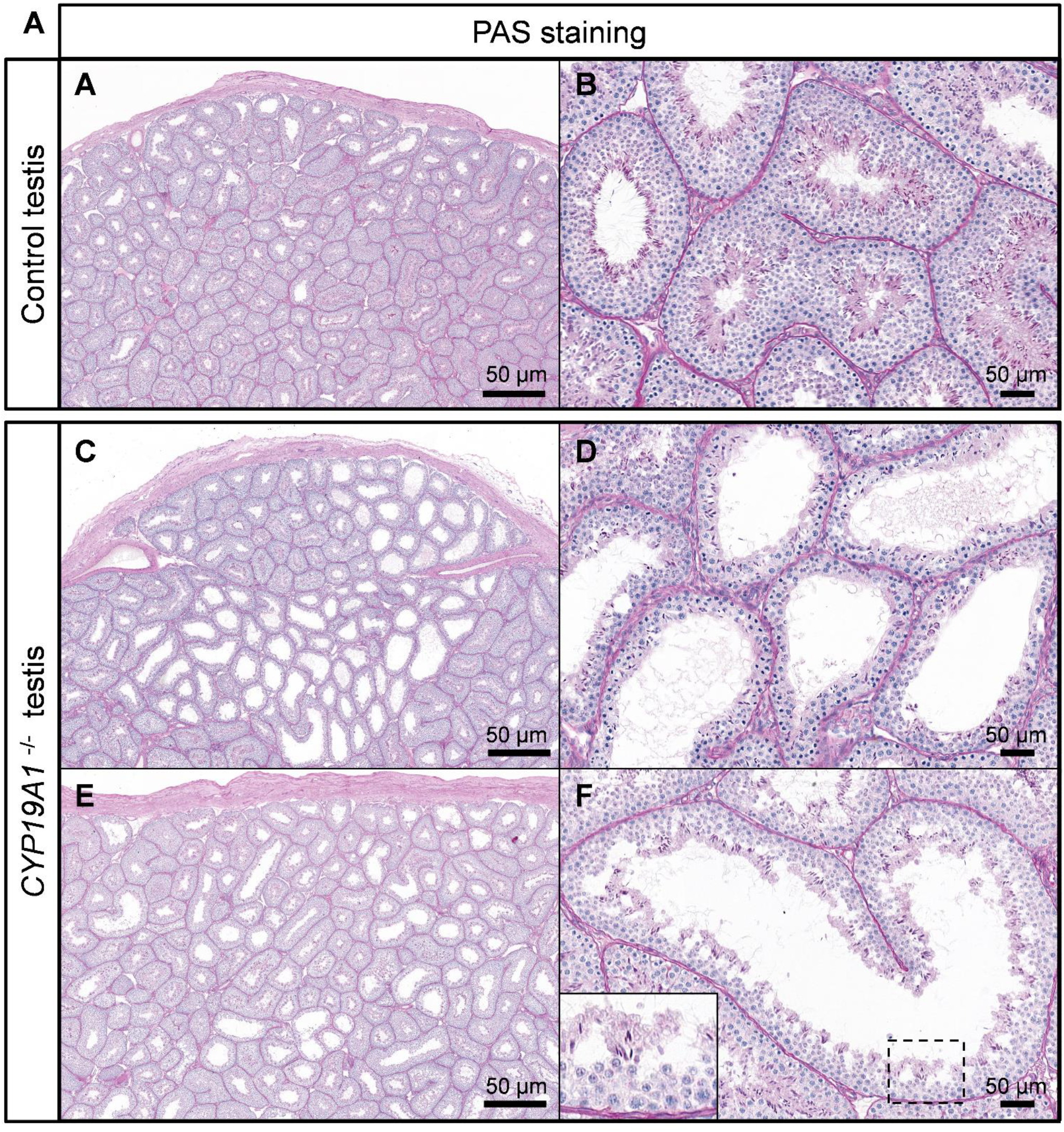
Spermatogenesis defects in absence of estrogen production. (**A-B**) PAS staining on control testis. (**C-F**) PAS staining on *CYP19A1*^−/−^ testis from two different rabbits (C-D and E-F). Males are 2 to 3 years old.

Molecular analyses confirmed a defect in the transition from spermatocytes to spermatids, with decreased mRNA levels of *SYCP3* (pValue = 0.057) and *PRM1* (Protamine 1) (pValue = 0.02) (Figure 5A and B). Consistent with spermatogenesis abnormalities, testis to body weight ratio was found to be significantly lower in *CYP19A1^−/−^* males compared to control ones (Figure 5C). In addition, total sperm number was estimated using the IVOS II CASA system (computer assisted sperm analysis), showing that the sperm count was significantly decreased in absence of estrogen synthesis (Figure 5D).

**Figure 5.**
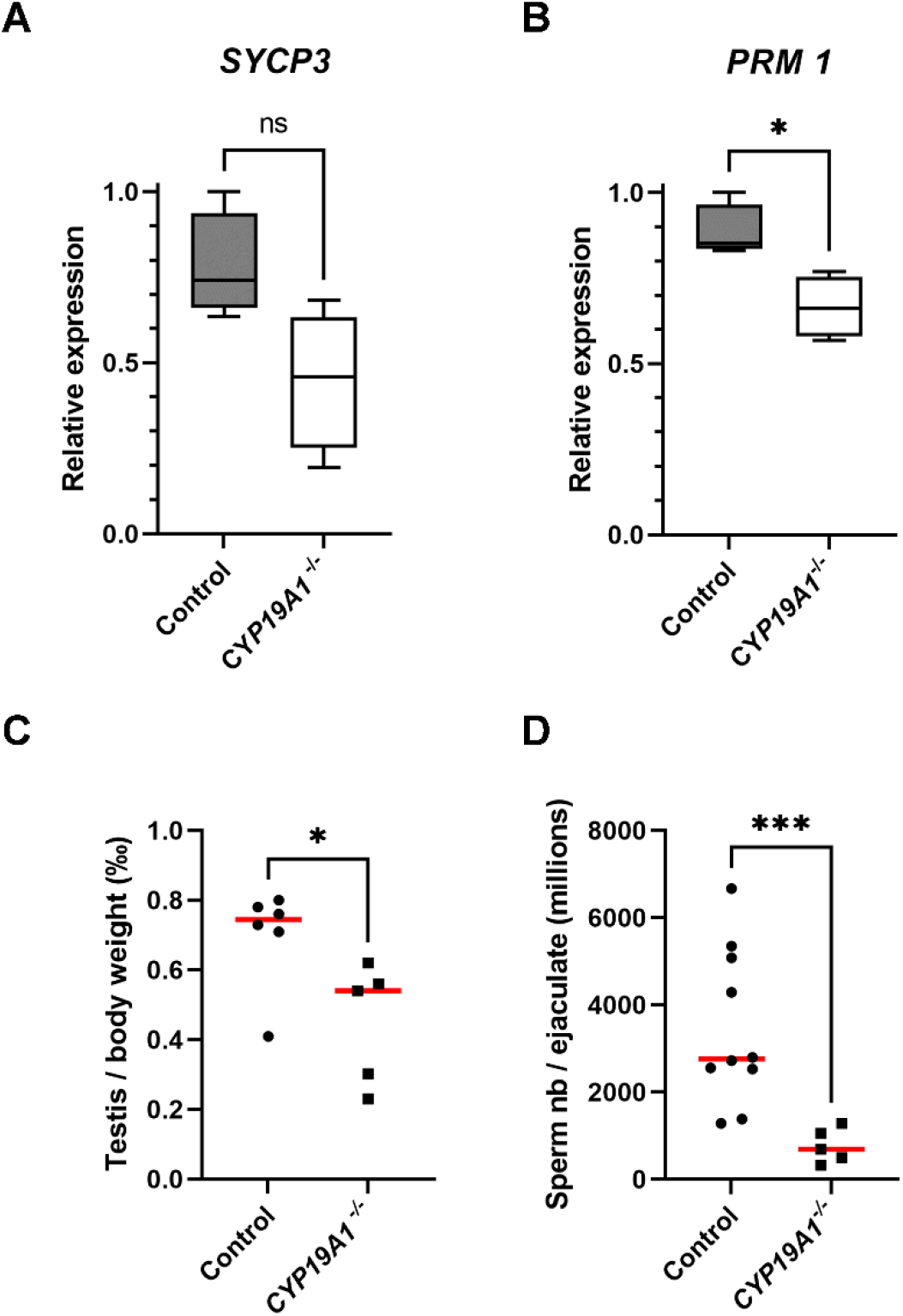
Hypo-spermatogenesis in *CYP19A1^−/−^* males. (**A-B**) RT-qPCR analyses of mRNA levels of *SYCP3* and *PRM1* in control and *CYP19A1*^−/−^ adult testis (n=5 for each genotype). (**C**) Testis on body weight ratio in control and *CYP19A1*^−/−^ rabbits. Control, n=6; *CYP19A1*^−/−^ n = 5. The median is shown in red. (**D**) Total sperm counts per million per ejaculated sample in control and *CYP19A1*^−/−^ rabbits. Control, n=10; *CYP19A1*^−/−^ n = 5. Dots represent the average of two successive semen collections per animal. For *CYP19A1*^−/−^ rabbits, three sets of two successive ejaculations were collected over a one-year interval. The median is shown in red. Mann-Whitney test: *pValue<0.05. ns: non significant.

### 3.5. Absence of testicular estrogens has no impact on germ cell DNA methylation

Since estrogen receptor over activation has been linked to epigenetic modifications [24], we were interested in the DNA methylation of germ cells. Nevertheless, immunofluorescence studies of the deposition of 5-methyl Cytosine (DNA methylation) or the 5-hydroxymethyl Cytosine (DNA hydroxymethylation, i.e. DNA demethylation) showed no difference between control and *CYP19A1*^−/−^ testes (Figure 6A). In addition, the DNA methylation rate of ejaculated sperm was determined by luminometric methylation assay (LUMA). The percentage of DNA methylation, around 70%, was found identical between sperm from control and mutant rabbits (Figure 6B), suggesting that if estrogen plays a role in epigenetics of the male gamete, this might not have been detected by global DNA methylation assessment.

**Figure 6.**
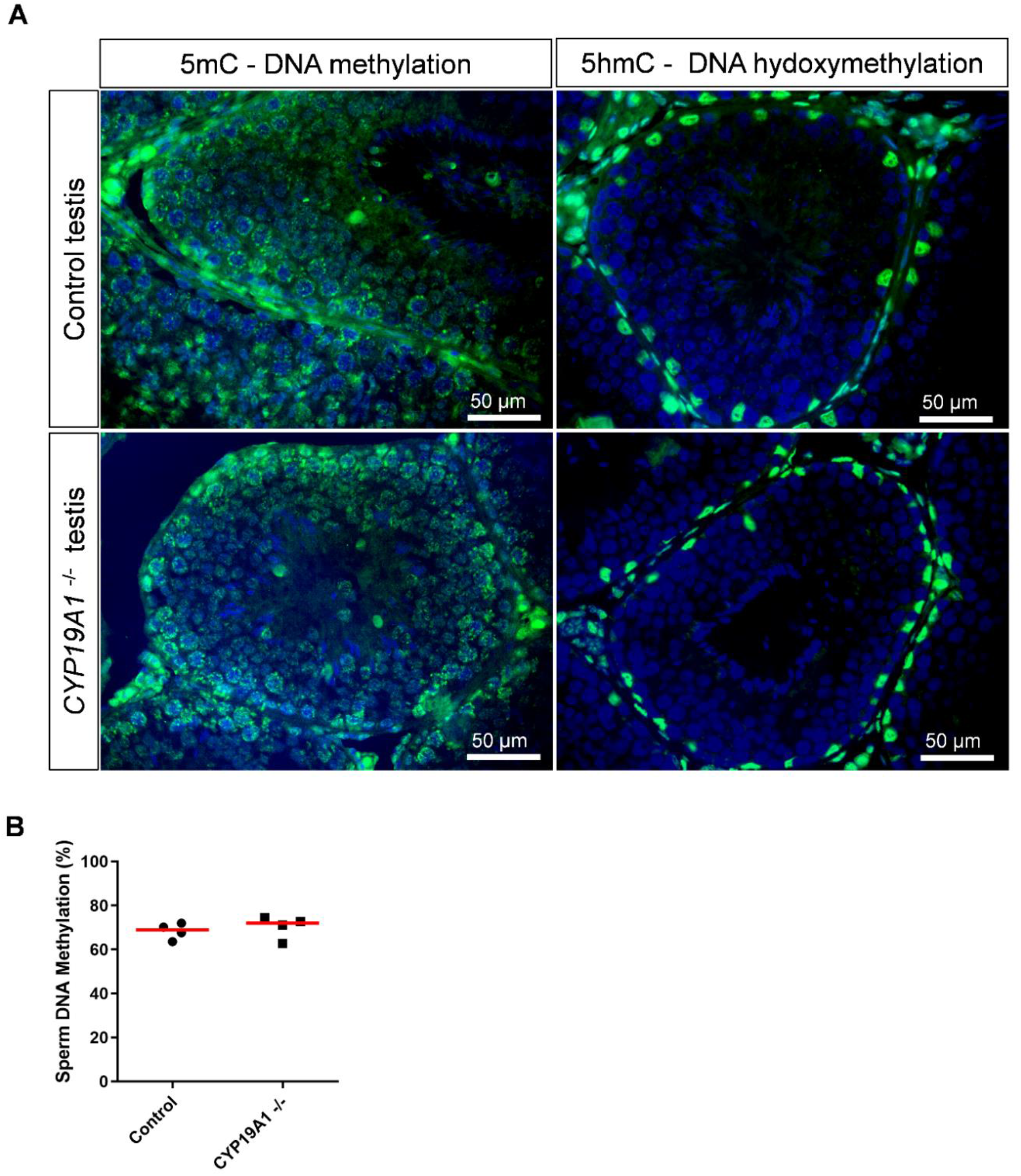
DNA methylation in the testis and sperm in absence of estrogen synthesis. (**A**) Immunodetection of 5mC and 5hmC in control and *CYP19A1*^−/−^ testis (green). Nuclei were stained in blue (DAPI). (**B**) Percentage of sperm DNA methylation from control and *CYP19A1^−/−^* male rabbits determined by Luminometric Methylation Assay (LUMA). The median is shown in red. Mann-Whitney test: nonsignificant.

### 3.6. Absence of testicular estrogen leads to sperm defects

In order to better understand the subfertility of *CYP19A1^−/−^* males, the sperm parameters of mutant and control rabbits were evaluated using the IVOS II CASA system on ejaculated sperm. Of the parameters assessed by this procedure, six were found statistically divergent between control and mutant sperm (Figure 7A-F). First, we noticed a decrease in total (from 90% to 55 %) and progressive (from 60% to 30%) sperm motility in mutant rabbits (Figure 7A and 7B). Second, the mutants displayed increased sperm malformations such as bent tails and Distal Midpiece Reflex curvatures (DMR) (Figure 7C and 7D). Finally, the spermatozoa from mutant animals retained more proximal and distal (Figure 7E and 7F) cytoplasmic droplets than the control sperm, suggesting an imperfect maturation of the gametes during their transit in the epididymis [42].

**Figure 7.**
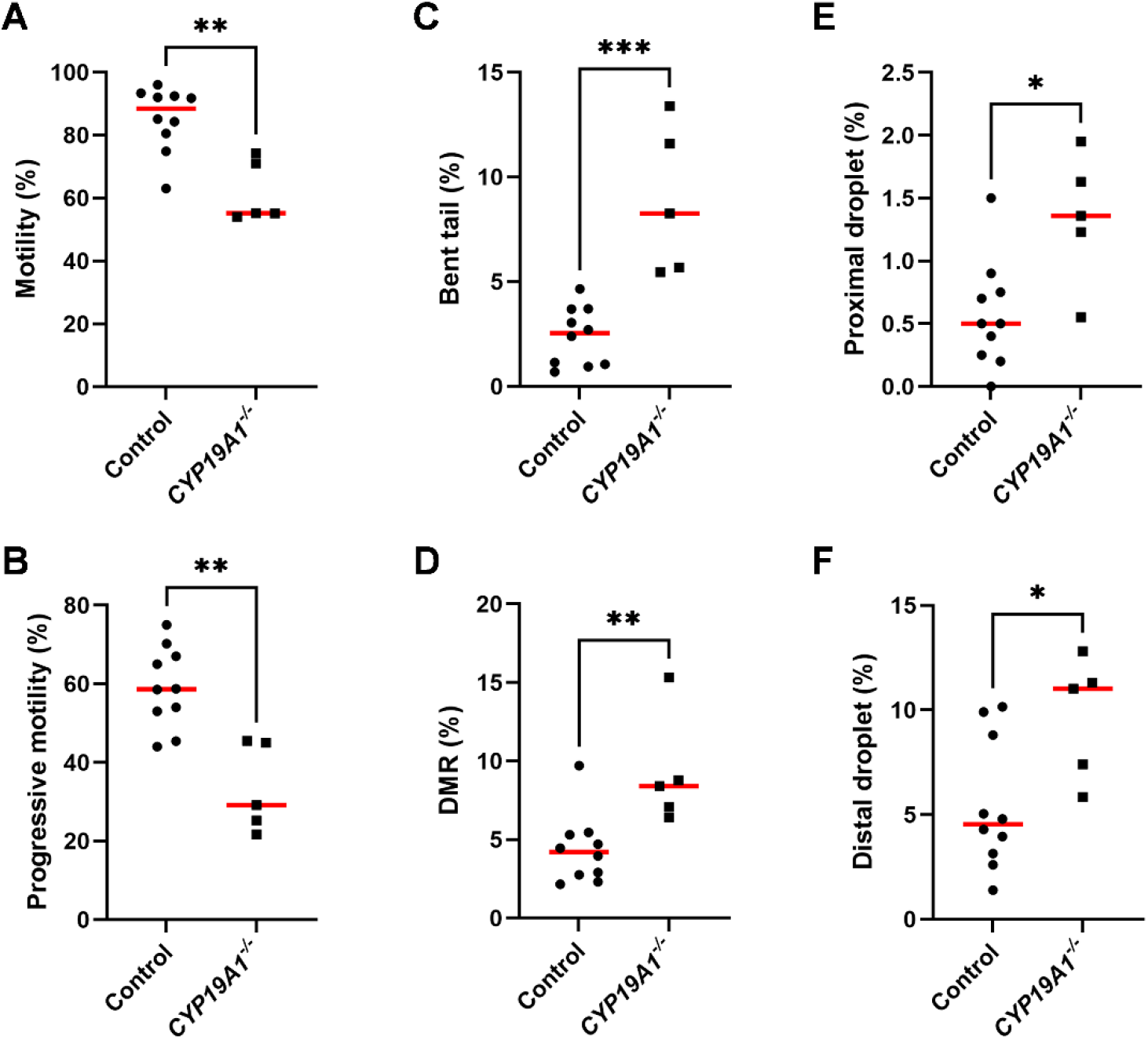
Spermatic parameters in control and *CYP19A1*^−/−^ rabbits. Motility and morphometric parameters of the sperm from control and *CYP19A1*^−/−^ rabbits were obtained by Computer Assisted Sperm Analysis. Percentages of (**A**) total motility and (**B**) progressive motility of the sperm were decreased in mutants. Percentages of (**C**) bent tails and (**D**) Distal Midpiece Reflex were increased in *CYP19A1*^−/−^ semen, together with (**E**) proximal droplets and (**F**) distal droplets. Dots represent the average of two successive semen collections per animal. For *CYP19A1*^−/−^ rabbits, three sets of two successive ejaculations were collected over a one-year interval. The median is shown in red. Control n= 10; *CYP19A1* ^−/−^ n= 5. Mann-Whitney test: * pValue<0.05; ** pValue<0.005; *** pValue<0.0005.

## 4. Discussion

In rabbits, the production of estrogen by the adult testes is strictly limited to the germ cells inside the seminiferous tubules: mainly to the meiotic germ cells (pachytene spermatocytes), but also slightly to the post-meiotic ones (round spermatids). While the location of *aromatase* expression differs according to the published studies, several of them are concordant on this point in rodents and in humans [9,14,15]. Thus, because of the blood-testis barrier established in the tubules, testicular estrogens should not pass through the general circulation in rabbits, but rather have a local function on the germ cells themselves, or on the male genital tract. Accordingly, 17β-Estradiol could be measured in the head and the tail of epididymis, showing that testicular estrogens diffuse into the fluid during the epididymal transit.

### 4.1. Testicular estrogens are involved in germ cells differentiation

In the testis, both *ESR1* and *ESR2* receptors were detected in round spermatids, suggesting that estrogens may have a role on post-meiotic germ cells. Accordingly, hypo-spermatogenesis has been observed in homozygous *CYP19A1^−/−^* mutant males, with some testicular lobules showing thinner seminiferous epithelium, with a lack of round spermatids, as described in the ArKO mouse model [31,32]. In the rat, overstimulation of ESR1 or ESR2 leads to spermiogenesis defects [22]. Thus, a lack or an excess of estrogens may impact spermatid differentiation. Additionally, epigenetic defects in the spermatids have been reported when estrogen receptors were overstimulated [23,24]. But, in this present study, we could not detect any changes in DNA methylation in the absence of estrogen. This may be related to the use of inappropriate methods, but estrogen might also not be involved in epigenetic reprograming in normal situations.

In humans, loss-of-function mutations affecting *CYP19A1*/*Aromatase* gene are very rare and poorly documented. In one reported case, the authors described abnormal skeletal growth and bone maturation in a male patient, which were associated with testicular hypoplasia and infertility [35]. The initiation of a replacement therapy by daily injection of estrogens restored the bone/skeletal phenotype, but had no effect on the testicles or fertility disorders. This last aspect could be linked to the fact that estrogens must be produced locally in the seminiferous tubules in order to be able to act on the differentiation of germ cells. In addition, a testicular biopsy, performed in this patient, revealed hypo-spermatogenesis and an arrest of germ differentiation, mainly at the level of primary spermatocytes [35]. This phenotype is close to that observed in rabbits, where round spermatids were rare in some tubules of the *CYP19A1*^−/−^ males, suggesting either that estrogens are necessary for their differentiation or maintenance, or that estrogens may be involved in the completion of meiosis. Interestingly, overexposure to estrogen or BPA has the same impact on spermatogenesis, with meiotic progression being defective and stopping at the pachytene stage [43]. In addition, in the female, the resumption of oocyte meiosis (with the first polar body extrusion) has been shown to be improved *in vitro* by the addition of estrogens to the medium [44]. In the present study, in rabbits deficient for aromatase, *SYCP3* and *PRM1* mRNAs levels were found to be decreased (pValue = 0.057 and 0.02 respectively) which could be an additional clue to consider the function of estrogens on meiotic cells. Further investigations involving high throughput transcriptome sequencing may highlight the potential implication of estrogens into meiosis in males.

### 4.2. Testicular estrogens are involved in sperm production, maturation and motility

As a consequence of the described defect in spermatogenesis, mild but significant testicular hypoplasia was observed in *CYP19A1*^−/−^ rabbits and the number of ejaculated spermatozoa decreased. These animals presented slight subfertility with conception difficulties (unsuccessful mating), as well as a decrease in the number of offspring. In addition of a decrease in sperm count, an increase in sperm abnormalities was observed. First, like in humans [34,35] and mice [29,31,32], sperm motility was affected, with a 50% reduction of the progressive motility in *CYP19A1*^−/−^ compared to control males. Then, related to the motility defects, increased percentages of flagellar abnormalities were noted, including bent tail and Distal Midpiece Reflex (DMR) which were doubled in mutants. In addition, proportion of sperm with cytoplasmic residual droplets was increased. These last phenotypes could be related to sperm maturation trouble in the epididymis, where sperm motility is acquired. In particular, the cytoplasmic droplet is expected to migrate caudally along the sperm during epididymal transit, and this droplet has been implicated in affecting some biochemical aspects of motility [45]. Some signaling pathways have been associated with sperm motility, and evidence suggests that sperm may have functional flagellar machinery that is activated during epididymal transit [45]. Indeed, in the epididymis, sperm undergo protein changes. As sperm are translationally silent, proteins appearing in them are thought to be synthesized by the epididymal epithelium and then incorporated to the sperm cells, thanks to exosomes for instance, called epididymosomes [46]. ESR1 and ESR2 receptors were both detected in the epididymis of male rabbits, mainly in the tail, where estrogens seem thus to exert their functions. In particular, ESR2 was found in the epithelial cells, which are supposed to secrete epididymosomes. Additional analyses on the transcriptomes and proteomes of mutant epididymides could provide clues to better understand how estrogen pathway dysfunctions impact sperm maturation and motility.

## 5. Conclusion

In the rabbit, testicular estrogens are produced by meiotic germ cells inside the seminiferous tubules. They play two main roles in promoting the fertility of the male gamete: (i) on germ cells and their progression in spermatogenesis; (ii) on the epididymis and indirectly on sperm maturation and motility acquisition. The phenotype of the *CYP19A1^−/−^* rabbits is very similar to the rare cases of *aromatase* mutations reported in humans, making the rabbit a relevant biomedical model for understanding and preventing male fertility.

## Author Contributions

Conceptualization, G.J. and M.P.; Methodology, A.D., E.D., M.A., A.A., F.G., H.J., E.S., G.J and M.P.; Validation, A.D., E.D., A.A, M.A., and H.J.; Formal Analysis, A.D., E.D., E.P. and M.P. Investigation, A.D., G.J and M.P.; Writing – Original Draft Preparation, M.P., E.P. and A.D; Writing – Review & Editing, H.J, G.J, E.P. and M.P.; Supervision, M.P.; Funding Acquisition, G.J. and E.P..

## Institutional Review Board Statement

The study was conducted according to the guidelines of Declaration of Helsinki, and approved by the French Ministry MENESR (accreditation number APAFIS#6775-2016091911407980 vI) following the recommendation given by the local committee for ethic in animal experimentation (COMETHEA, Jouy-en-Josas, France). All researchers working directly with the animals possessed an animal experimentation license delivered by the French veterinary services.

## Data Availability Statement

Not applicable.

## Acknowledgments

The authors would like to thank Patrice Congar, Gwendoline Morin and all the staff of the facility (SAAJ, INRAE, Jouy-en-Josas) for the care of the rabbits and semen collection, and Julie Rivière and Marthe Vilotte (UMR GABI, INRAE, Jouy-en-Josas) for their assistance on the histological platform (@Bridge platform) and for the access to the virtual slide scanner.

## References

1. Guiguen, Y.; Baroiller, J.F.; Ricordel, M.J.; Iseki, K.; Mcmeel, O.M.; Martin, S.A.; Fostier, A. Involvement of Estrogens in the Process of Sex Differentiation in Two Fish Species: The Rainbow Trout (Oncorhynchus Mykiss) and a Tilapia (Oreochromis Niloticus). Mol Reprod Dev 1999, 54, 154–162, doi:10.1002/(SICI)1098-2795(199910)54:2<154∷AID-MRD7>3.0.CO;2-5.

2. Pieau, C.; Dorizzi, M.; Richard-Mercier, N. Temperature-Dependent Sex Determination and Gonadal Differentiation in Reptiles. Cell Mol Life Sci 1999, 55, 887–900, doi:10.1007/s000180050342.

3. Elbrecht, A.; Smith, R.G. Aromatase Enzyme Activity and Sex Determination in Chickens. Science 1992, 255, 467–470, doi:10.1126/science.1734525.

4. Wade, J.; Arnold, A.P. Functional Testicular Tissue Does Not Masculinize Development of the Zebra Finch Song System. Proc Natl Acad Sci U S A 1996, 93, 5264–5268, doi:10.1073/pnas.93.11.5264.

5. Vaillant, S.; Dorizzi, M.; Pieau, C.; Richard-Mercier, N. Sex Reversal and Aromatase in Chicken. J Exp Zool 2001, 290, 727–740, doi:10.1002/jez.1123.

6. Jost, A. A New Look at the Mechanisms Controlling Sex Differentiation in Mammals. Johns Hopkins Med J 1972, 130, 38–53.

7. Hess, R.A.; Sharpe, R.M.; Hinton, B.T. Estrogens and Development of the Rete Testis, Efferent Ductules, Epididymis and Vas Deferens. Differentiation 2021, 118, 41–71, doi:10.1016/j.diff.2020.11.004.

8. Lambard, S.; Silandre, D.; Delalande, C.; Denis-Galeraud, I.; Bourguiba, S.; Carreau, S. Aromatase in Testis: Expression and Role in Male Reproduction. J Steroid Biochem Mol Biol 2005, 95, 63–69, doi:10.1016/j.jsbmb.2005.04.020.

9. Levallet, J.; Bilinska, B.; Mittre, H.; Genissel, C.; Fresnel, J.; Carreau, S. Expression and Immunolocalization of Functional Cytochrome P450 Aromatase in Mature Rat Testicular Cells1. Biology of Reproduction 1998, 58, 919–926, doi:10.1095/biolreprod58.4.919.

10. Fraczek, B.; Kotula-Balak, M.; Wojtusiak, A.; Pierściński, A.; Bilińska, B. Cytochrome P450 Aromatase in the Testis of Immature and Mature Pigs. Reprod Biol 2001, 1, 51–59.

11. Sipahutar, H.; Sourdaine, P.; Moslemi, S.; Plainfossé, B.; Séralini, G.-E. Immunolocalization of Aromatase in Stallion Leydig Cells and Seminiferous Tubules. J Histochem Cytochem 2003, 51, 311–318, doi:10.1177/002215540305100306.

12. Payne, A.H.; Kelch, R.P.; Musich, S.S.; Halpern, M.E. Intratesticular Site of Aromatization in the Human. J Clin Endocrinol Metab 1976, 42, 1081–1087, doi:10.1210/jcem-42-6-1081.

13. Papadopoulos, V.; Carreau, S.; Szerman-Joly, E.; Drosdowsky, M.A.; Dehennin, L.; Scholler, R. Rat Testis 17β-Estradiol: Identification by Gas Chromatography-Mass Spectrometry and Age Related Cellular Distribution. Journal of Steroid Biochemistry 1986, 24, 1211–1216, doi:10.1016/0022-4731(86)90385-7.

14. Nitta, H.; Bunick, D.; Hess, R.A.; Janulis, L.; Newton, S.C.; Millette, C.F.; Osawa, Y.; Shizuta, Y.; Toda, K.; Bahr, J.M. Germ Cells of the Mouse Testis Express P450 Aromatase. Endocrinology 1993, 132, 1396–1401, doi:10.1210/endo.132.3.8440194.

15. Lambard, S.; Galeraud-Denis, I.; Saunders, P.T.K.; Carreau, S. Human Immature Germ Cells and Ejaculated Spermatozoa Contain Aromatase and Oestrogen Receptors. J Mol Endocrinol 2004, 32, 279–289, doi:10.1677/jme.0.0320279.

16. Rago, V.; Aquila, S.; Panza, R.; Carpino, A. Cytochrome P450arom, Androgen and Estrogen Receptors in Pig Sperm. Reprod Biol Endocrinol 2007, 5, 23, doi:10.1186/1477-7827-5-23.

17. Dostalova, P.; Zatecka, E.; Dvorakova-Hortova, K. Of Oestrogens and Sperm: A Review of the Roles of Oestrogens and Oestrogen Receptors in Male Reproduction. Int J Mol Sci 2017, 18, E904, doi:10.3390/ijms18050904.

18. Pelletier, G.; El-Alfy, M. Immunocytochemical Localization of Estrogen Receptors Alpha and Beta in the Human Reproductive Organs. J Clin Endocrinol Metab 2000, 85, 4835–4840, doi:10.1210/jcem.85.12.7029.

19. Fietz, D.; Ratzenböck, C.; Hartmann, K.; Raabe, O.; Kliesch, S.; Weidner, W.; Klug, J.; Bergmann, M. Expression Pattern of Estrogen Receptors α and β and G-Protein-Coupled Estrogen Receptor 1 in the Human Testis. Histochem Cell Biol 2014, 142, 421–432, doi:10.1007/s00418-014-1216-z.

20. Cavaco, J.E.B.; Laurentino, S.S.; Barros, A.; Sousa, M.; Socorro, S. Estrogen Receptors Alpha and Beta in Human Testis: Both Isoforms Are Expressed. Syst Biol Reprod Med 2009, 55, 137–144, doi:10.3109/19396360902855733.

21. Hirata, S.; Shoda, T.; Kato, J.; Hoshi, K. Isoform/Variant MRNAs for Sex Steroid Hormone Receptors in Humans. Trends Endocrinol Metab 2003, 14, 124–129, doi:10.1016/s1043-2760(03)00028-6.

22. Dumasia, K.; Kumar, A.; Deshpande, S.; Sonawane, S.; Balasinor, N.H. Differential Roles of Estrogen Receptors, ESR1 and ESR2, in Adult Rat Spermatogenesis. Mol Cell Endocrinol 2016, 428, 89–100, doi:10.1016/j.mce.2016.03.024.

23. Dumasia, K.; Kumar, A.; Deshpande, S.; Balasinor, N.H. Estrogen, through Estrogen Receptor 1, Regulates Histone Modifications and Chromatin Remodeling during Spermatogenesis in Adult Rats. Epigenetics 2017, 12, 953–963, doi:10.1080/15592294.2017.1382786.

24. Dumasia, K.; Kumar, A.; Deshpande, S.; Balasinor, N.H. Estrogen Signaling, through Estrogen Receptor β, Regulates DNA Methylation and Its Machinery in Male Germ Line in Adult Rats. Epigenetics 2017, 12, 476–483, doi:10.1080/15592294.2017.1309489.

25. Krege, J.H.; Hodgin, J.B.; Couse, J.F.; Enmark, E.; Warner, M.; Mahler, J.F.; Sar, M.; Korach, K.S.; Gustafsson, J.A.; Smithies, O. Generation and Reproductive Phenotypes of Mice Lacking Estrogen Receptor Beta. Proc Natl Acad Sci U S A 1998, 95, 15677–15682, doi:10.1073/pnas.95.26.15677.

26. Antal, M.C.; Krust, A.; Chambon, P.; Mark, M. Sterility and Absence of Histopathological Defects in Nonreproductive Organs of a Mouse ERbeta-Null Mutant. Proc Natl Acad Sci U S A 2008, 105, 2433–2438, doi:10.1073/pnas.0712029105.

27. Otto, C.; Fuchs, I.; Kauselmann, G.; Kern, H.; Zevnik, B.; Andreasen, P.; Schwarz, G.; Altmann, H.; Klewer, M.; Schoor, M.; et al. GPR30 Does Not Mediate Estrogenic Responses in Reproductive Organs in Mice. Biol Reprod 2009, 80, 34–41, doi:10.1095/biolreprod.108.071175.

28. Eddy, E.M.; Washburn, T.F.; Bunch, D.O.; Goulding, E.H.; Gladen, B.C.; Lubahn, D.B.; Korach, K.S. Targeted Disruption of the Estrogen Receptor Gene in Male Mice Causes Alteration of Spermatogenesis and Infertility. Endocrinology 1996, 137, 4796–4805, doi:10.1210/endo.137.11.8895349.

29. Joseph, A.; Hess, R.A.; Schaeffer, D.J.; Ko, C.; Hudgin-Spivey, S.; Chambon, P.; Shur, B.D. Absence of Estrogen Receptor Alpha Leads to Physiological Alterations in the Mouse Epididymis and Consequent Defects in Sperm Function. Biol Reprod 2010, 82, 948–957, doi:10.1095/biolreprod.109.079889.

30. Joseph, A.; Shur, B.D.; Ko, C.; Chambon, P.; Hess, R.A. Epididymal Hypo-Osmolality Induces Abnormal Sperm Morphology and Function in the Estrogen Receptor Alpha Knockout Mouse. Biol Reprod 2010, 82, 958–967, doi:10.1095/biolreprod.109.080366.

31. Robertson, K.M.; O’Donnell, L.; Jones, M.E.E.; Meachem, S.J.; Boon, W.C.; Fisher, C.R.; Graves, K.H.; McLachlan, R.I.; Simpson, E.R. Impairment of Spermatogenesis in Mice Lacking a Functional Aromatase (Cyp 19) Gene. Proceedings of the National Academy of Sciences 1999, 96, 7986–7991, doi:10.1073/pnas.96.14.7986.

32. Robertson, K.M.; Simpson, E.R.; Lacham-Kaplan, O.; Jones, M.E. e. Characterization of the Fertility of Male Aromatase Knockout Mice. Journal of Andrology 2001, 22, 825–830, doi:10.1002/j.1939-4640.2001.tb02587.x.

33. Haverfield, J.T.; Ham, S.; Brown, K.A.; Simpson, E.R.; Meachem, S.J. Teasing out the Role of Aromatase in the Healthy and Diseased Testis. Spermatogenesis 2011, 1, 240, doi:10.4161/spmg.1.3.18037.

34. Herrmann, B.L.; Saller, B.; Janssen, O.E.; Gocke, P.; Bockisch, A.; Sperling, H.; Mann, K.; Broecker, M. Impact of Estrogen Replacement Therapy in a Male with Congenital Aromatase Deficiency Caused by a Novel Mutation in the CYP19 Gene. J Clin Endocrinol Metab 2002, 87, 5476–5484, doi:10.1210/jc.2002-020498.

35. Carani, C.; Qin, K.; Simoni, M.; Faustini-Fustini, M.; Serpente, S.; Boyd, J.; Korach, K.S.; Simpson, E.R. Effect of Testosterone and Estradiol in a Man with Aromatase Deficiency. N Engl J Med 1997, 337, 91–95, doi:10.1056/NEJM199707103370204.

36. Jolivet, G.; Daniel-Carlier, N.; Harscoët, E.; Airaud, E.; Dewaele, A.; Pierson, C.; Giton, F.; Boulanger, L.; Daniel, N.; Mandon-Pépin, B.; et al. Fetal Estrogens Are Not Involved in Sex Determination But Critical for Early Ovarian Differentiation in Rabbits. Endocrinology 2022, 163, bqab210, doi:10.1210/endocr/bqab210.

37. Hellemans, J.; Mortier, G.; De Paepe, A.; Speleman, F.; Vandesompele, J. QBase Relative Quantification Framework and Software for Management and Automated Analysis of Real-Time Quantitative PCR Data. Genome Biology 2007, 8, R19, doi:10.1186/gb-2007-8-2-r19.

38. Giton, F.; Sirab, N.; Franck, G.; Gervais, M.; Schmidlin, F.; Ali, T.; Allory, Y.; de la Taille, A.; Vacherot, F.; Loric, S.; et al. Evidence of Estrone-Sulfate Uptake Modification in Young and Middle-Aged Rat Prostate. J Steroid Biochem Mol Biol 2015, 152, 89–100, doi:10.1016/j.jsbmb.2015.05.002.

39. Devillers, M.M.; Petit, F.; Cluzet, V.; François, C.M.; Giton, F.; Garrel, G.; Cohen-Tannoudji, J.; Guigon, C.J. FSH Inhibits AMH to Support Ovarian Estradiol Synthesis in Infantile Mice. J Endocrinol 2019, 240, 215–228, doi:10.1530/JOE-18-0313.

40. Perrier, J.-P.; Sellem, E.; Prézelin, A.; Gasselin, M.; Jouneau, L.; Piumi, F.; Al Adhami, H.; Weber, M.; Fritz, S.; Boichard, D.; et al. A Multi-Scale Analysis of Bull Sperm Methylome Revealed Both Species Peculiarities and Conserved Tissue-Specific Features. BMC Genomics 2018, 19, 404, doi:10.1186/s12864-018-4764-0.

41. Karimi, M.; Johansson, S.; Stach, D.; Corcoran, M.; Grandér, D.; Schalling, M.; Bakalkin, G.; Lyko, F.; Larsson, C.; Ekström, T.J. LUMA (LUminometric Methylation Assay)--a High Throughput Method to the Analysis of Genomic DNA Methylation. Exp Cell Res 2006, 312, 1989–1995, doi:10.1016/j.yexcr.2006.03.006.

42. Cooper, T.G. Cytoplasmic Droplets: The Good, the Bad or Just Confusing? Human Reproduction 2005, 20, 9–11, doi:10.1093/humrep/deh555.

43. Liu, C.; Duan, W.; Li, R.; Xu, S.; Zhang, L.; Chen, C.; He, M.; Lu, Y.; Wu, H.; Pi, H.; et al. Exposure to Bisphenol A Disrupts Meiotic Progression during Spermatogenesis in Adult Rats through Estrogen-like Activity. Cell Death Dis 2013, 4, e676, doi:10.1038/cddis.2013.203.

44. Chi, H.; Cao, Z. Effect of Oestrogen on Mouse Follicle Growth and Meiotic Resumption. Zygote 2022, 30, 330–337, doi:10.1017/S0967199421000708.

45. Gervasi, M.G.; Visconti, P.E. Molecular Changes and Signaling Events Occurring in Sperm during Epididymal Maturation. 2018, 30.

46. Sullivan, R.; Saez, F. Epididymosomes, Prostasomes, and Liposomes: Their Roles in Mammalian Male Reproductive Physiology. Reproduction 2013, 146, R21–35, doi:10.1530/REP-13-0058.

